# Gradient based refinement of CryoET tilt series alignment improves tomogram contrast and structure resolution

**DOI:** 10.64898/2026.01.16.699989

**Authors:** Muyuan Chen

## Abstract

Cryogenic electron tomography (CryoET) provides 3D views of vitrified cellular samples, and protein structures can be determined from the tomograms by averaging many copies of the same protein computationally. However, the resolution of these averaged structures, particularly for smaller proteins, is often constrained by the precision of tilt-series alignment. In this study, we introduce a gradient descent-based approach to refine alignment parameters, enhancing the contrast in tomograms of the sample regions. This refinement not only improves contrast but also yields higher-resolution protein structures derived from the same particle populations.

## Introduction

While traditional structure biology studies mainly target biochemically purified proteins, only inside native cellular environment can we observe the full structural complexity of the macromolecular machines. With the technological development of Cryo-FIB-SEM and CryoET, as well as the advance of data processing methods, it has become possible to determine protein structures *in situ* at near atomic resolution^1^. However, current high resolution *in situ* structure studies are often limited to large protein complexes, often 1MDa or larger, such as ribosomes^2,3^, microtubules^4^, or respiratory complexes^5,6^.

CryoET provides 3D views of the vitrified samples by tilting the stage at a series of orientations inside the microscope, providing a stack of micrographs, or a tilt series. The tilt series can then be reconstructed into a tomogram, where densities of individual proteins can be identified. Hundreds of thousands of copies of the same protein are aligned and averaged to overcome the low signal to noise ratio (SNR) in raw micrographs and produce 3D structures of the target protein at high resolution.

One of the key challenges in CryoET based structure determination is to achieve accurate tilt series alignment. Even with thin, purified protein samples that have gold fiducials that can be used as landmarks for computational alignment, it has been shown that the tilt series alignment is still highly inaccurate and limits the resolution of subtomogram averaging^7^. The lamella generated by Cryo-FIB-SEM makes the problem even more challenging, since gold fiducials or other high contrast landmarks are generally absent in the lamella, and the complexity of various cellular features further limits the accuracy of patch tracking based alignment method.

Currently, the main solution to overcome the misalignment of tilt series is reference based sub-tilt refinement during the subtomogram averaging process^7–9^. While the naming and implementation of the protocol varies in different software packages, the basic concept is similar. In addition to searching the 3D Euler angle of each subtomogram particle, the algorithm also locally refine the translation/rotation of the sub-tilt series that contribute to the 3D subtomogram, using projections of averaged 3D structures as references. The sub-tilt refinement has shown great improvement in the resolution achieved from subtomogram averaging, making it possible to subnanometer or near atomic resolution structures for large complexes like ribosomes. However, the success of sub-tilt refinement is limited by the SNR of individual sub-tilt images, which are much noisier than the 3D subtomograms that are used for the 3D Euler angle search. As a result, it is more challenging to determine structures of smaller proteins inside cells using CryoET. While it is possible to use the information from multiple particles in a surrounding region to boost the SNR for the sub-tilt alignment ^9^, it is still difficult to achieve results comparable to the case of large complexes. The multi-particle refinement approach is also situational because not every particle has enough neighbors in a small enough surrounding region that have good 3D reference and can be used for the alignment.

## Method

To address this, here we present a gradient descent based algorithm that refines the alignment of tilt series and maximizes the contrast of the sample region. Using a deep learning package ^10^, we implement the tilt series alignment and tomogram reconstruction steps in a differentiable model. This makes it possible to automatically trace the gradient of the contrast in the final tomogram with respect to the alignment parameters of the tilt series. Through iterative refinement using a gradient based optimizer, we can achieve tilt series alignment parameters that produce tomograms with higher sample contrast. Using the refined tilt series alignment parameters, we can also produce better 3D particles that can achieve higher resolution through iterative subtomogram alignment and averaging.

To refine the tilt series alignment, we start from an existing set of alignment parameters obtained from classical landmark based alignment or patch tracking. The tilt series is downsampled so that the Nyquist frequency corresponds to 50Å resolution, and we divide the tomogram region into multiple cubic tiles along the X-Y axis. Each tile is independently reconstructed by inserting the corresponding 2D slices in the Fourier using trilinear interpolation. The 3D tiles are normalized so that average is zero and the standard deviation of top and bottom slab is one. For each tile, we compute the slice-by-slice standard deviation of each 2D image along the z axis (Figure 1A-C). To optimize the alignment parameters, we use the average standard deviation of the central z slices across all tiles as the loss function. That is, we aim to maximize the contrast of the central slab of the tomogram where the biological sample resides, against the top and bottom of the tomogram where there is mostly noise. The reconstruction protocol is implemented using the Jax library, which performs the automatic differentiation of the tomogram contrast with respect to the input tilt series alignment parameters. An Adam optimizer is used to refine the alignment parameters iteratively and maximize the contrast.

**Figure 1.**
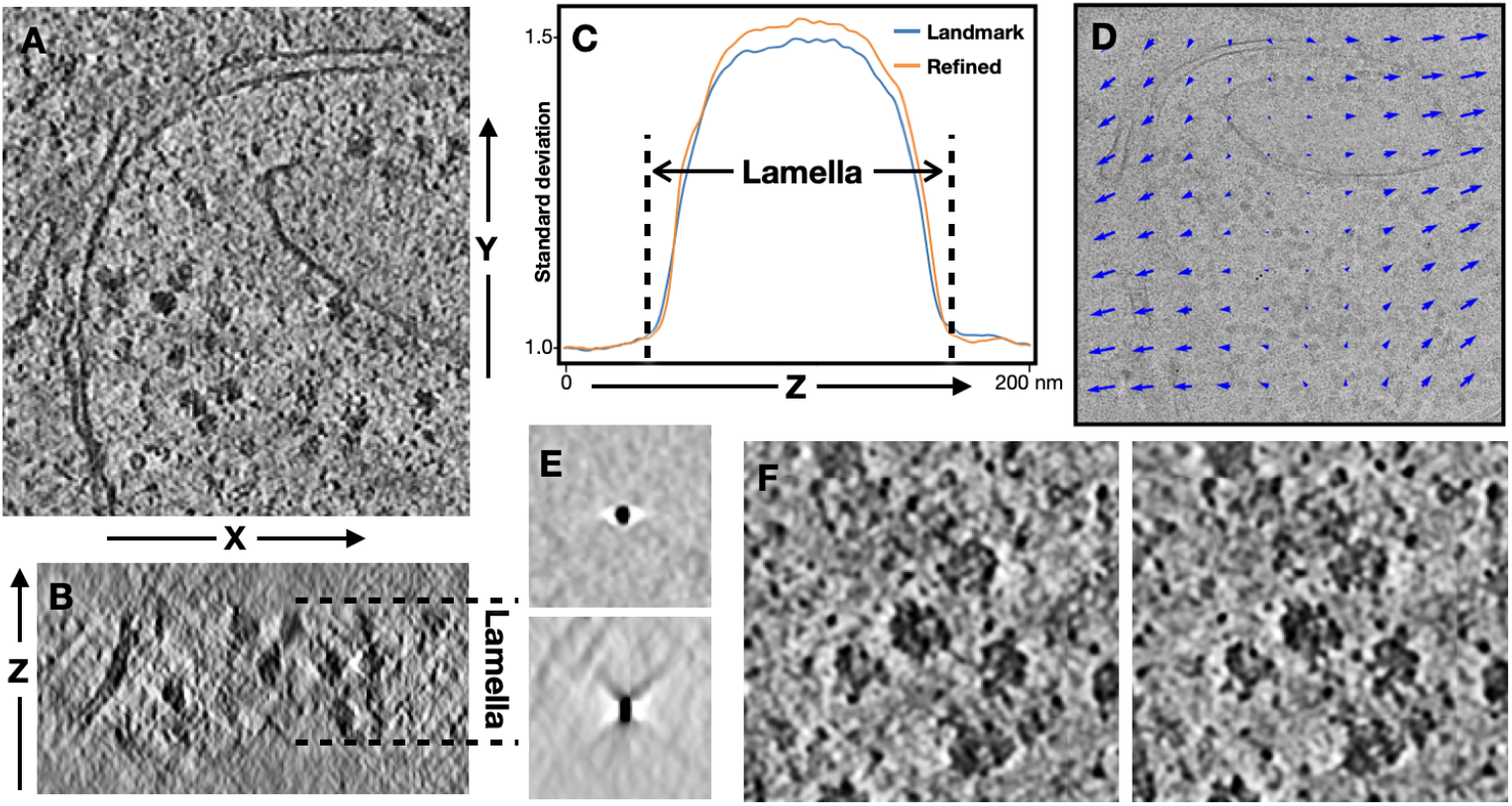
Refinement of tilt series alignment parameters. (A) Slice view of one tomogram, viewed from the X-Y plane. (B) The tomogram viewed from the X-Z plane, with dashed lines showing the region with lamella. (C) Standard deviation of each X-Y slice of the tomogram, plotted along the Z axis. Blue - tomogram reconstructed using landmark based alignment; orange - tomogram reconstructed using refined parameters with local motion considered. (D) Trajectories of local motion on one of the near center tilt images obtained from the refinement. Length of arrows is exaggerated. (E) Top and side view of one of the high contrast fiducials in the tomogram, reconstructed using the standard landmark based alignment parameters. (F) Zoomed in view of a tomogram region reconstructed using landmark based alignment (left) and refined parameters (right).

A few notable adjustments are implemented to improve the result of the refinement. First, an inverted leaky ReLU function is applied to the reconstructed 3D tiles after normalization^11^. This essentially upweights the negative contrast, which corresponds to densities of biomolecules in the tomogram, in the optimization. Second, the voxel values are squared before calculating standard deviation. This further upweights high contrast voxels that are more likely protein densities instead of noise. Third, during the iterative optimization, we randomly set 5% of the alignment parameters back to the original ones from the classical tilt series alignment results. This is similar to the drop out process during neural network training that stabilizes the optimization process^12^. Finally, the central slab for optimization is by default defined as the middle 40% along the z axis. When coordinates of target particles are provided, we alternatively only use the range of z slices that have particles, so we avoid the impact of high contrast objects (e.g. crystalized ice) above and below the sample.

In addition to the X-Y translation of the tilt images, the algorithm can optionally consider the local drift within each image of the tilt series. For simplification, here we model the local motion as first order polynomials (Figure 1D). That is, the drift of each tile is a function of the X-Y coordinate at the center of that tile. During the refinement, the algorithm directly optimizes the coefficients of the polynomial model, and the additional translation of each tile is calculated from the coefficients. To extract 3D particles after the refinement, we multiply the particle coordinates with the coefficient to obtain the location of the sub-tilt particles on each micrograph.

## Results

To evaluate the result of the tilt series alignment, we apply the method to tilt series of a public CryoET dataset of green algae cell lamellae (EMPIAR-11830)^13^. We start from 5 tilt series that contain cytosolic ribosomes. In addition to cellular features, these tilt series have multiple small and high contrast particles on the surface of the lamella, which can be used as fiducials for the classical landmark based tilt series alignment ^8^. The alignment yields high quality tomograms, evident by the undistorted side view of the fiducials (Figure 1E). From the y-z plane view of the tomogram, the boundary between the biological features within the lamella and the vacuum above and below is clearly visible. After the gradient based refinement, the contrast of the lamella content is increased numerically, even though the reconstructed tomograms do not show obvious differences in structure features visually (Figure 1F).

2271 ribosome particles were selected from the 5 tomograms (Figure 2A), and an existing ribosome structure, lowpass filtered and phase randomized to 30Å, was used as the initial reference for the iterative subtomogram refinement. Particle extraction and subtomogram refinement were performed independently following identical protocols for tilt series aligned using landmark based alignment and gradient based refinement methods. For each set of particles, 3 rounds of subtomogram alignment and 3 rounds of sub-tilt alignment were performed. To avoid potential bias from half-set splitting, masking and other factors, here we report resolution by calculating the Fourier shell correlation (FSC) between the subtomogram average and an existing high resolution ribosome structure, and report the resolution by FSC=0.5 cutoff.

**Figure 2.**
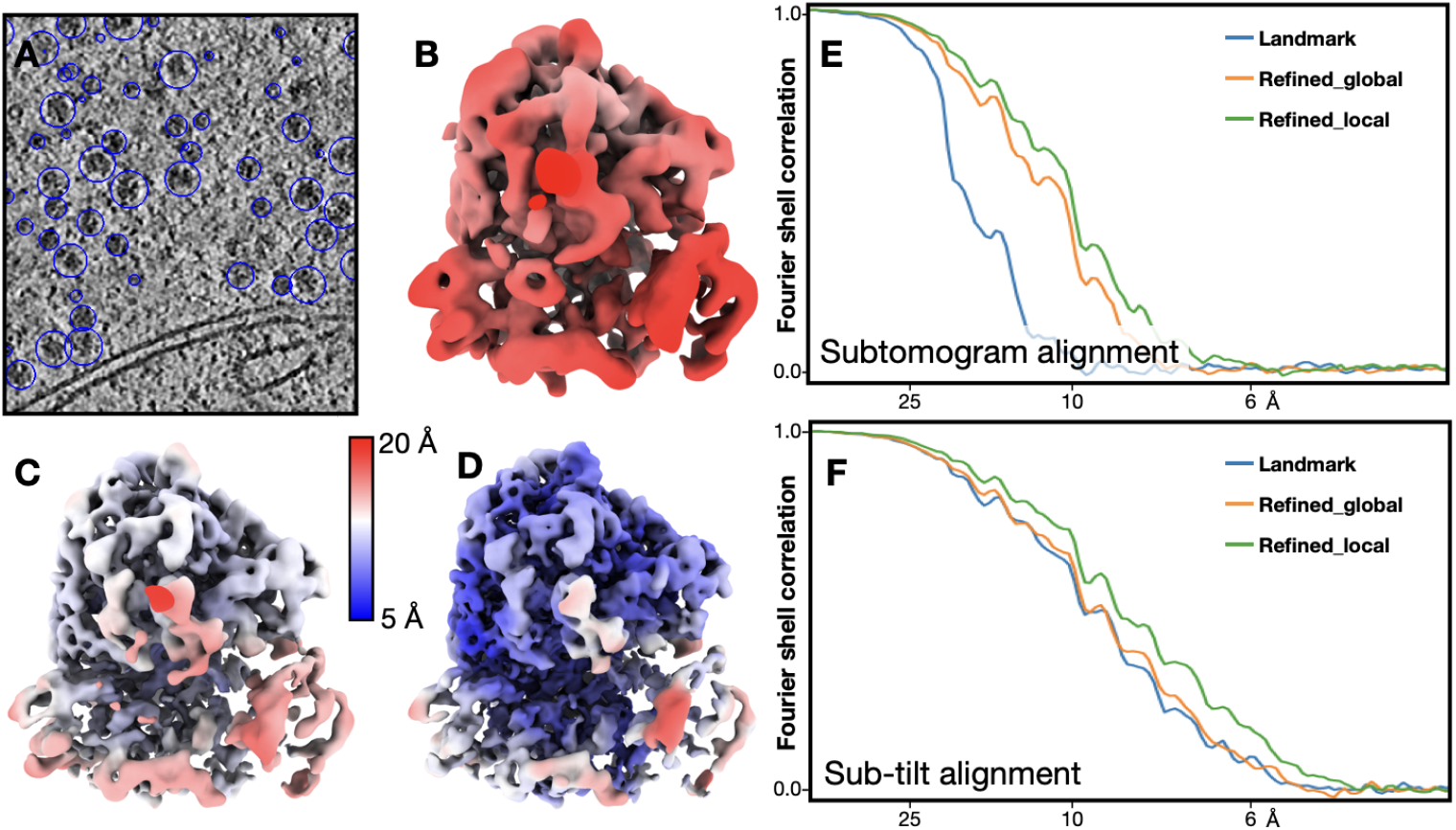
Subtomogram averaging of ribosomes. (A) Slice view of a tomogram with ribosome particles circled. (B) Averaged structure of ribosomes using tilt series alignment parameters from landmark based alignment, after only subtomogram alignment. The map is colored by local resolution. (C) Averaged structure of ribosomes using refined tilt series alignment parameters including local motion, after only subtomogram alignment. (D) Averaged structure of ribosomes using refined tilt series alignment parameters including local motion, after sub-tilt alignment. (E) Fourier shell correlation curves of ribosome refinement, using different tilt series alignment parameters, after only subtomogram alignment. (F) Fourier shell correlation curves of ribosome refinement, after sub-tilt alignment.

With only subtomogram alignment and using the tilt series alignment parameters, the subtomogram average only achieved 16.7 Å resolution (Figure 2B). The resolution improved to 9.6 Å after sub-tilt alignment. Using tilt series alignment parameters from the gradient based refinement, but without considering the local drift, the averaged structure can achieve 10.2 Å resolution with only subtomogram alignment. After the sub-tilt alignment, the resolution again arrived at 9.5 Å, and the structure obtained is virtually the same as the one from classical tilt series alignment. Finally, using refined tilt series alignment parameters that also considers local motion, the resolution is 9.8 Å after subtomogram alignment (Figure 2C), and 8.5 Å after sub-tilt alignment (Figure 2D), a slight improvement compared to the classical method. Overall, in the case of ribosomes, while the refinement of tilt series alignment using the new method showed meaningful improvement of subtomogram average resolution, most of the improvement can be similarly achieved using sub-tilt refinement (Figure 2E-F).

Next, we move on to test the method on a more challenging example, the RubisCOs inside the chloroplast of the same green algae dataset. Here, we selected 12,395 RubisCO particles from 3 tomograms (Figure 3A). Compared to the ribosomes, this is a much less ideal case for subtomogram averaging. RubisCOs are much smaller than ribosomes, only ∼15% in molecular weight. The dataset includes only 3 tomograms, which means the oscillation of the CTF may not be fully flattened through averaging. Finally, even though the RubisCOs are densely packed in the tomograms, during the sub-tilt refinement, we only use 15 nearest neighbors for the alignment, mimicking a less crowded situation closer to the environment that most proteins are in outside the condensate. Similar to the ribosome example, we used an existing RubisCO structure (EMD-22401)^14^, lowpass filtered to 30Å, as reference, and performed 3 rounds of subtomogram alignment followed by 3 rounds of sub-tilt alignment. D4 symmetry was applied throughout the refinement.

**Figure 3.**
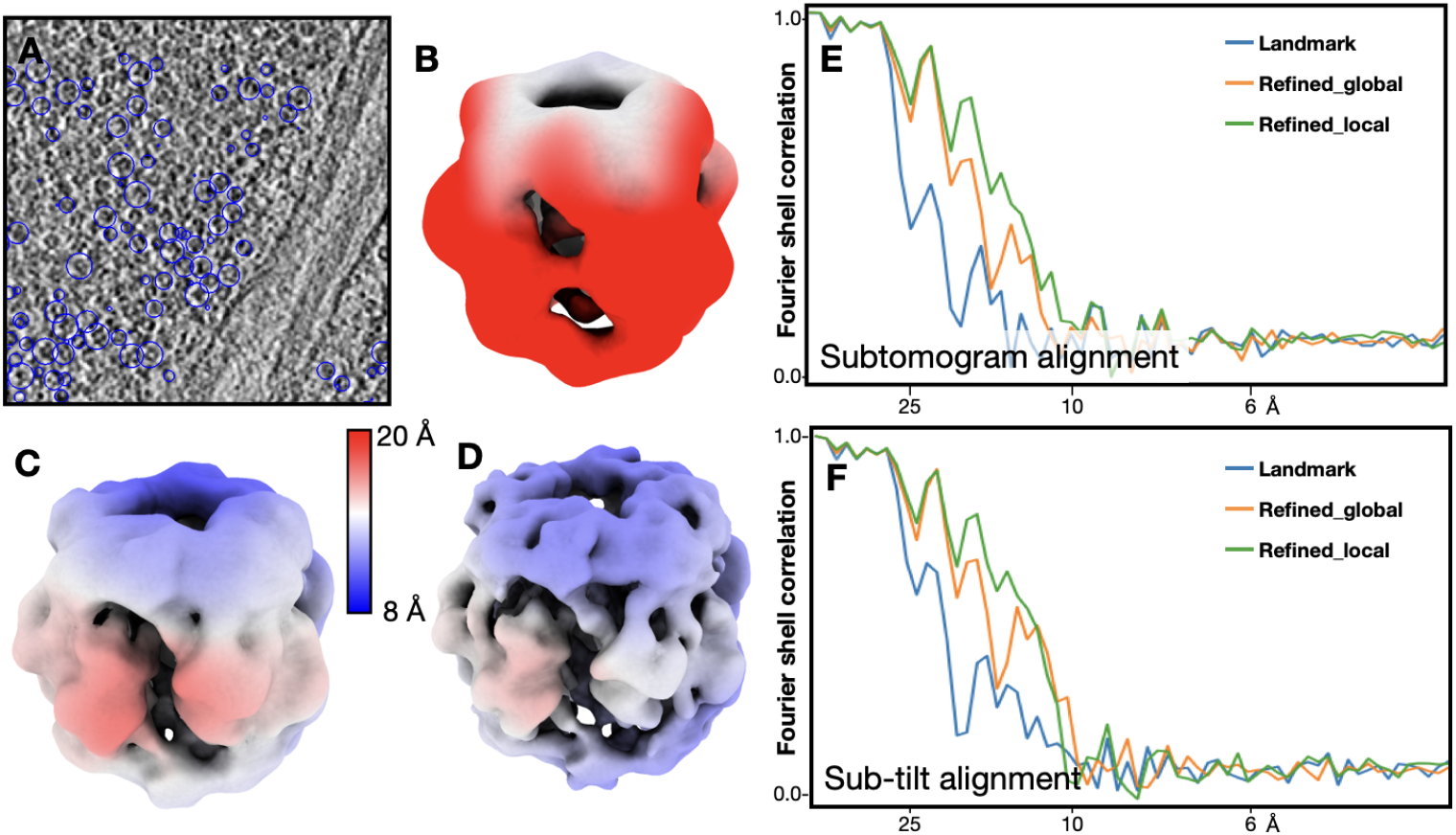
Subtomogram averaging of RubisCOs. (A) Slice view of a tomogram with RubisCOs particles circled. (B) Averaged structure of RubisCOs using tilt series alignment parameters from landmark based alignment, after only subtomogram alignment. The map is colored by local resolution. (C) Averaged structure of RubisCOs using refined tilt series alignment parameters including local motion, after only subtomogram alignment. (D) Averaged structure of RubisCOs using refined tilt series alignment parameters including local motion, after sub-tilt alignment. (E) Fourier shell correlation curves of RubisCO refinement, using different tilt series alignment parameters, after only subtomogram alignment. (F) Fourier shell correlation curves of RubisCO refinement, after sub-tilt alignment.

With alignment parameters from the classical tilt series alignment routine, the resolution of subtomogram average is 36.5 Å after only subtomogram alignment (Figure 3B), and 26.1 Å after sub-tilt alignment. With refined tilt series alignment parameters without local motion, the resolution reached 24.3 Å after subtomogram alignment, and 20.3 Å after sub-tilt refinement. If we consider local motion in tilt series refinement, the resolution of RubisCO reached 19.2 Å after subtomogram alignment (Figure 3C), which is improved to 16.6 Å after sub-tilt refinement (Figure 3D). In this example, because the size of the protein limits the performance of sub-tilt alignment, the improvement of the tilt series alignment refinement becomes dominant, and structures with higher resolution can be obtained with the exact same particles (Figure 3E-F).

## Discussion

In sum, here we introduced an algorithm for tilt series alignment refinement that uses tomogram contrast at the sample region as the target loss function. Using the refined alignment, higher resolution can be achieved through subtomogram averaging without reference based alignment of sub-tilt series. While the impact of the additional tilt series refinement is relatively limited for large complexes (e.g. ribosomes), the method showed a clear improvement for smaller proteins, where the sub-tilt alignment cannot fully correct the misalignment due to the low SNR per particle. This method will potentially expand the types of proteins we can target for subtomogram averaging from the lamella, and allow us to move a step forward toward CryoET based visual proteomics.

In addition to the simple test cases shown here, the refined tilt series alignment also shows improvement in structure resolution in large datasets. For example, using the full dataset, we are able to obtain RubisCO structures at 6.5Å resolution^15^, compared to the 7.5Å resolution structure from the original publication^16^. Notably, extensive particle classification and selection was required to achieve the 7.5Å structure from the original paper (only 6% of original picks were used), while all particles that were initially selected from tomograms were used in our RubisCO structure. While some of this can be attributed to better particle picking that avoids non-particle features, and some may be caused by better heterogeneity analysis, it is possible that tilt series with sub-optimal alignment contribute to a large portion of the “bad particles”. Having better tilt series alignment from the beginning can potentially lead to having more “good particles” in the averaged structures, and a larger dataset also leaves more room for comprehensive structure heterogeneity analysis afterwards.

Despite the success, there are clear rooms of improvement for this method. First, the loss function we use simply uses standard deviation to represent the contrast in the tomograms. While it is a reasonable approximation, the standard deviation can also be affected by the noise and other artifacts in CryoET data. Ideally, in place of the lowpass filter currently used, some feature aware denoising method can be applied to the tomogram before calculating the loss function^17,18^, so we are measuring the actual contrast of cellular features in the tomogram instead of everything combined. While multiple CryoET denoising methods already exist, their implementation is currently limited by the GPU memory, since it is challenging to fit a neural network denoising model and the current tilt series refinement protocol into a single GPU.

Second, the polynomial form that we used to describe the local motion within the lamella is naive and may not be a good representation of the actual movement. With enough CryoET data and the corresponding local motion trajectory computed from sub-tilt refinement of ribosomes, it should be possible to find a good function form for the deformation of the lamella. With the good function form, the refinement just needs to fit a few parameters using information from the current tilt series to obtain a realistic pattern of local motion.

The tilt series alignment refinement method described in this paper is distributed with the EMAN2 software package, and can be accessed using the e2tomogram_refine.py program.

## Acknowledgement

This research is supported by NIH grant R01GM150905.

